# Genomic analyses correspond with deep persistence of peoples of Blackfoot Confederacy from glacial times

**DOI:** 10.1101/2023.09.25.559300

**Authors:** Dorothy First Rider, Annabel Crop Eared Wolf, John Murray, Alida de Flamingh, Andre Luiz Campelo dos Santos, François Lanoë, Maria N. Zedeño, Michael DeGiorgio, John Lindo, Ripan S. Malhi

## Abstract

Mutually beneficial partnerships between genomics researchers and North American Indigenous Nations are rare. Here, we present one such partnership that provides insight into the peopling of the Americas and furnishes a new line of evidence that can be used to further treaty and Indigenous rights. We show that the genomics of sampled individuals from the Blackfoot Confederacy belong to a previously undescribed ancient lineage that diverged from other genomic lineages in the Americas in Late Pleistocene times. Using multiple complementary forms of knowledge, we provide a scenario for Blackfoot population history that fits with oral tradition and provides a plausible model for the evolutionary process of the peopling of the Americas.

**One-Sentence Summary:** Previously unknown genomic lineage in North America revealed in present-day Indigenous community and historic ancestors.

---

Studies of human genomes have aided historical research in the Americas by providing rich information about demographic events and population histories of Indigenous peoples, including the initial peopling of the continents. The ability to study genomes of Ancestors in the Americas through paleogenomics has greatly increased the power and resolution at which we can infer past events and processes (*1*). However, few genomic studies have been completed with populations in North America, which could be the most informative about the initial peopling process. Those that have been completed in North America have identified Indigenous Ancestors with previously undescribed genomic lineages that evolved in the Late Pleistocene, prior to the split of two lineages (called the “Northern Native American (NNA)” or “ANC-B” and “Southern Native American (SNA)” or “ANC-A” lineages) from which all present-day Indigenous populations in the double continent that have been sampled derive much, if not all, their ancestry. Specifically, the lineage termed “Ancient Beringian” was ascribed to a genome in an Ancestor who lived 11,500 years ago at *Xaasaa Na’* (Upward Sun River) and named *Xach’itee’aanenh t’eede gaay* (USR1) by the local Healy Lake Village Council in Alaska. An Ancestor who lived 9,500 years ago at what is now called Trail Creek Caves on the Seward Peninsula, Alaska also belongs to the Ancient Beringian lineage (*2, 3*). Additionally, another Ancestor, under the stewardship of Stswecem’c Xgat’tem First Nation, who lived in what is now called British Columbia, belongs to a distinct genomic lineage that predates the NNA-SNA split, but postdates the split from Ancient Beringians on the Americas’ genomic timeline. This ancestor was identified at Big Bar Lake near the Frasier River and lived 5,600 years ago (*3, 4*). Thus, these previous studies of North American Indigenous Ancestors have successfully helped to identify previously unknown genomic diversity. However, the ancient lineages identified in these studies have not been observed in Indigenous peoples of the Americas living today. Research in Mesoamerica and South America suggests that certain sampled populations (e.g., Mixe) have at least partial ancestry in present-day Indigenous groups from unknown genomic lineages in the Americas, possibly dating as far back as 25,000 years ago (*3, 5, 6*)

In addition to providing information about the population histories of Indigenous peoples in the Americas, genomic studies have the potential to further Indigenous communities’ social, educational, legal, and political goals. Equitable partnerships between scientists and these communities are essential for advancing the fragmentary genomic knowledge of ancient and present-day Indigenous populations (*7*–*9*). This study presents results from such a mutually beneficial partnership among the Blood (Kainai) First Nation of Alberta, the Blackfeet Tribal Historic Preservation Office, and scientists (Indigenous and non-Indigenous) from various academic institutions.

The Blood (Kainai) First Nation and the Blackfeet Tribe are two of four geopolitical divisions of the Blackfoot Confederacy (Figure 1). Ancestors of the Blackfoot were mobile, specialized bison hunters who inhabited the Rocky Mountain Front and adjacent Northwestern Plains. Since the establishment of reservations in Canada and the United States, member tribes of the Blackfoot Confederacy have confronted biases about their origin and antiquity in land claims and water rights litigation and in the advancement and protection of their treaty and Indigenous rights (*10*–*12*). Yet, their long-term presence in the region can be traced, both archaeologically and through oral tradition, to the end of the last glaciation (*13, 14*).

**Figure 1.**
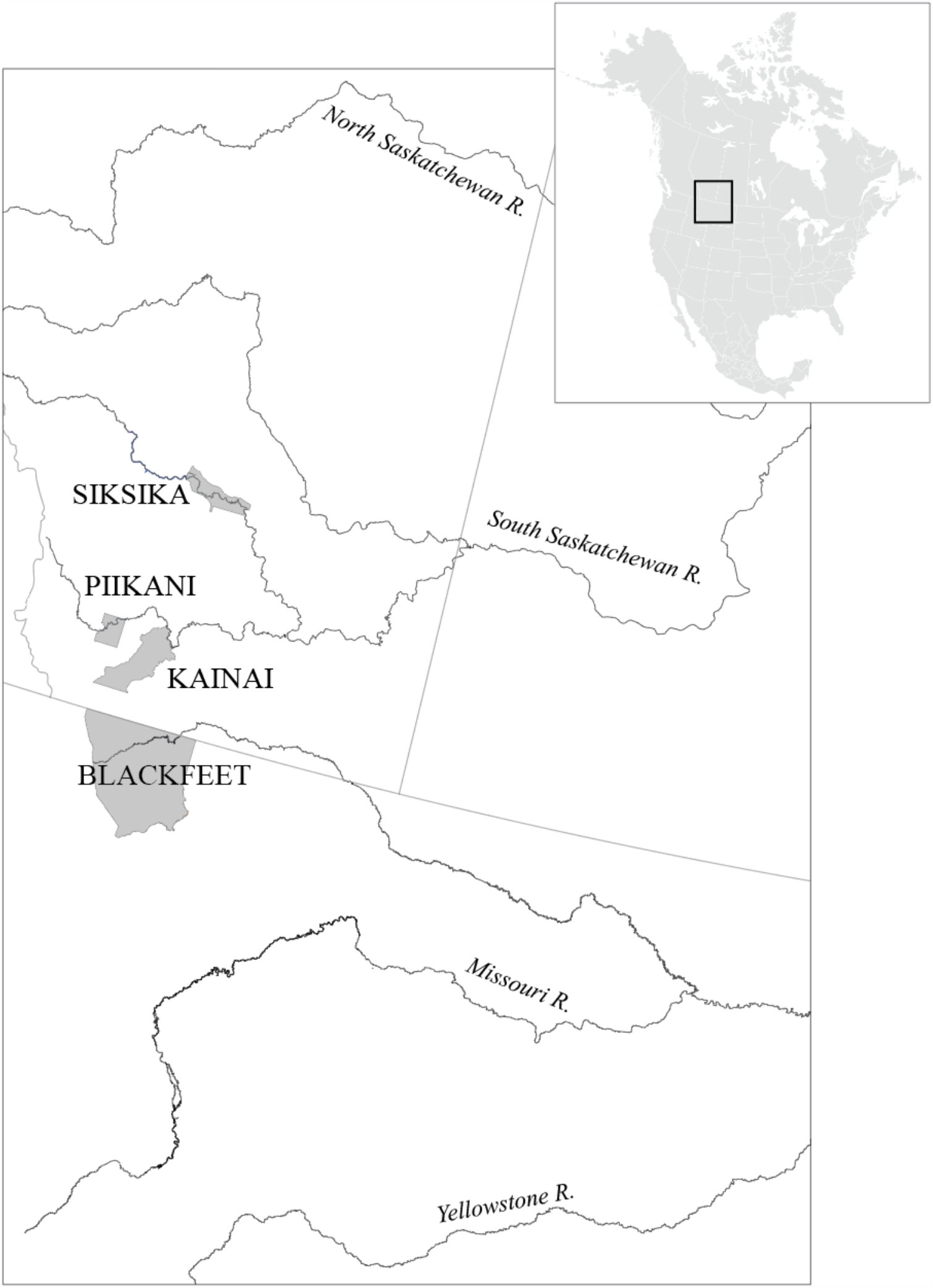
Map of Blackfoot traditional territory and the current geopolitical divisions of the Blackfoot Confederacy including Siksika, Pikani, Kainai and Blackfeet Nations.

Thus, the objectives of this study were not only to advance scientific knowledge about Indigenous genomic lineages that can provide insight into the peopling of the Americas, but also to provide the Blackfoot with an independent line of evidence for evaluating purported ancestral relationships with other North American groups. Specifically, the study provides insight into potential genetic affinities with Algonquian speakers and establishing claims of antiquity in the continent to assist with furthering treaty and Indigenous rights. Consequently, this study aimed at understanding genetic relationships between historical Ancestors and living individuals from Indigenous communities, modeling the population history and divergence of the Blackfoot genomic lineage, and mapping genetic relationships between these and other Native North American populations for which genomic data are available.

## Results

### Community engagement

Genomic research that is culturally responsive and works with communities for respectful co-production of knowledge is mutually beneficial and remodels scientific practices that are non-extractive (*7, 15*). To this end, in 2018 the Blood (Kainai) First Nation of Alberta, Canada and the Laboratory of Molecular Anthropology at the University of Illinois Urbana-Champaign signed a Memorandum of Understanding that presented agreed upon protocols and expectations for a study to analyze genomic data derived from historical Ancestors and contemporary individuals. The Blackfeet Tribal Historic Preservation Office (BTHPO), in conjunction with the Smithsonian Institution’s Repatriation Office, arranged for the sampling of historical Ancestors on behalf of the Blood First Nation. Archaeologists from the Bureau of Applied Research in Anthropology at The University of Arizona facilitated the partners’ interactions as part of the Blackfoot Early Origins Program, which is a multi-year and multi-project collaboration with member tribes of the Blackfoot Confederacy.

### Sampling and data generation

We completed low-coverage, whole genome sequencing on three historical Ancestors identified as Blood or Blackfoot from the Smithsonian Institute and four historical Ancestors curated at the Blackfeet Tribal Historic Preservation Office. DNA preservation in the samples was generally poor, with coverage ranging from 0.0027X to 0.0155X. However, we were able to infer genetic sex and mitochondrial DNA haplogroup for six of the seven historical Ancestors (Table 1). Two of the samples from historical Ancestors that exhibited the best DNA preservation in the set were re-sequenced to obtain coverages of 0.113X and 0.187X. DNA damage patterns showed nucleotide base-pair changes characteristic of ancient DNA, e.g., deamination of cytosine to uracil (Figure S1). Six present-day tribal members of the Blood or Blackfoot were also whole genome sequenced to a mean of 31X.

**Table 1.**
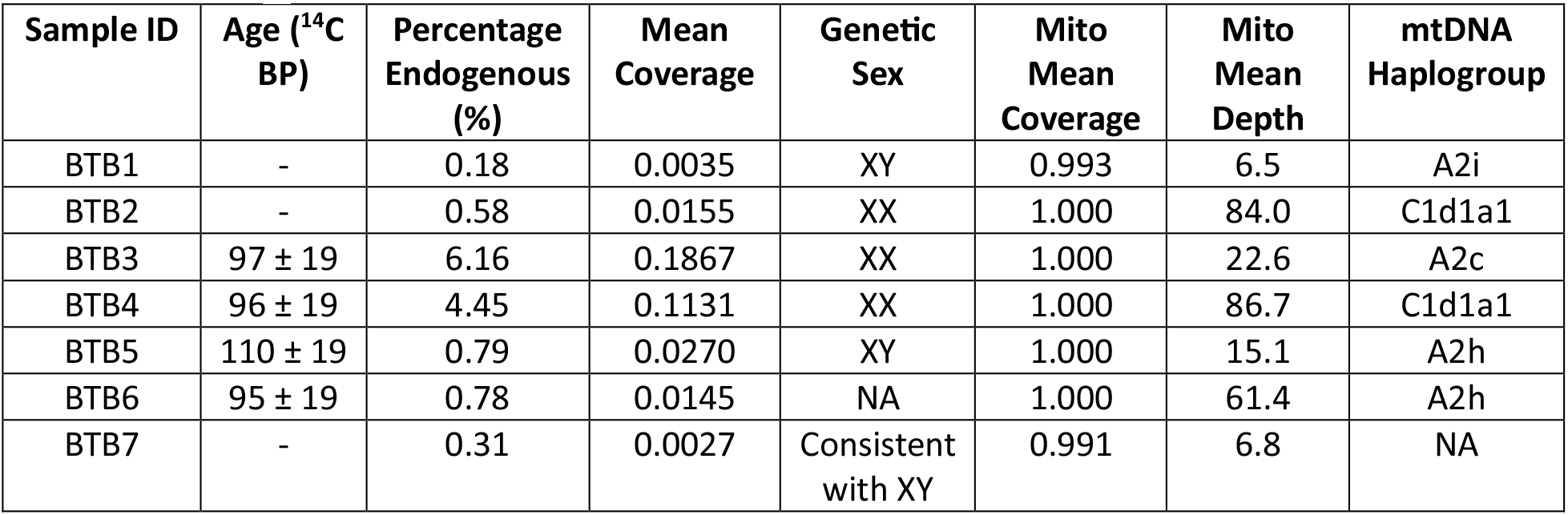
Samples from historic ancestors analyzed in this study with AMS date, percentage endogenous human DNA, mean coverage, genetic sex, mean mitochondrial DNA coverage, mean mitochondrial depth and mitochondrial DNA haplogroup reported.

| Sample ID | Age (<sup>14</sup>C BP) | Percentage Endogenous (%) | Mean Coverage | Genetic Sex | Mito Mean Coverage | Mito Mean Depth | mtDNA Haplogroup |
| --- | --- | --- | --- | --- | --- | --- | --- |
| BTB1 | - | 0.18 | 0.0035 | XY | 0.993 | 6.5 | A2i |
| BTB2 | - | 0.58 | 0.0155 | XX | 1.000 | 84.0 | C1d1a1 |
| BTB3 | 97 ± 19 | 6.16 | 0.1867 | XX | 1.000 | 22.6 | A2c |
| BTB4 | 96 ± 19 | 4.45 | 0.1131 | XX | 1.000 | 86.7 | C1d1a1 |
| BTB5 | 110 ± 19 | 0.79 | 0.0270 | XY | 1.000 | 15.1 | A2h |
| BTB6 | 95 ± 19 | 0.78 | 0.0145 | NA | 1.000 | 61.4 | A2h |
| BTB7 | - | 0.31 | 0.0027 | Consistent with XY | 0.991 | 6.8 | NA |

Radiocarbon assay of samples from the historical Ancestors produced ages ranging between 95 and 110 ^14^C BP (Table 1, Table S2). In all four cases, the reversal and plateau of the calibration curve at that time produces discontinuous calendar ages of 1685-1732 and 1805-1917 AD. The latter range is more likely given that scaffolds and associated surface remains would probably not have preserved for several centuries before collection. This time range is contemporaneous with an increase in Euromerican-Blackfoot interactions from the apogee of the fur trade era (ca. 1800-1860), introduction of infectious diseases such as smallpox, the Blackfoot War (1870), beginning of the reservation era (1873), and starvation winter (1883-84).

### Genomic structure and diversity in North America

To understand the relationship between the ancient and present-day individuals from the Blood and Blackfoot, as well as their relationship to other Indigenous populations from the Americas, we performed principal component analysis (PCA). The reference populations, including Indigenous populations, derived from the Simons Genome Diversity Project and ancient individuals from previously published studies (Table S1). The PCA contains genomic data from 41 individuals. Individuals from Europe cluster in the top right corner of the plot and individuals from Mesoamerica and South America are located at the bottom of the plot. The ancient individuals of the Blood/Blackfoot cluster together with present-day representatives of the Blood/Blackfoot and Cree groups in the plot of PC1 and PC2 (Figure 2A), but are more distant to Cree in additional PCA plots (Figure S2). The present-day Blood/Blackfoot exhibit a cline in the plot between ancient individuals from North America and individuals from Europe based on variation in ancestry among the present-day participants of the Blood/Blackfoot. Next, we inferred the relationship between the ancient Blood/Blackfoot, the present-day Blood/Blackfoot, and worldwide populations using maximum likelihood trees produced by *TreeMix*. The ancient Blood/Blackfoot and present-day Blood/Blackfoot lie on the same branch equal distance to branches with other groups from North America and South America (Figure 2B). This result is complemented by the *ADMIXTURE* analysis (with lowest cross-validation value at six ancestral components) that shows the ancient individuals of the Blood/Blackfoot exhibiting nearly all their ancestry belonging to the yellow component, with present-day Blood/Blackfoot and ancient North American individuals also demonstrating a large fraction of ancestry to the yellow component (Figure 2C). We also assessed the genomic relationship for groups in the Americas based on qpGraph analysis (Figure 3A), for which the best fitting model with no migration events exhibits a topology with ancient Blood/Blackfoot closest to the present-day Blood/Blackfoot. Lastly, *D*-statistics demonstrate an excess of allele sharing between the ancient Blood/Blackfoot and present-day Blood/Blackfoot (Figure 3B). In addition, ancient and present-day Blood/Blackfoot are equally distant to Indigenous populations in North America (apart from Cree due to high levels of European admixture) and South America (Figures 3 and S4).

**Figure 2.**
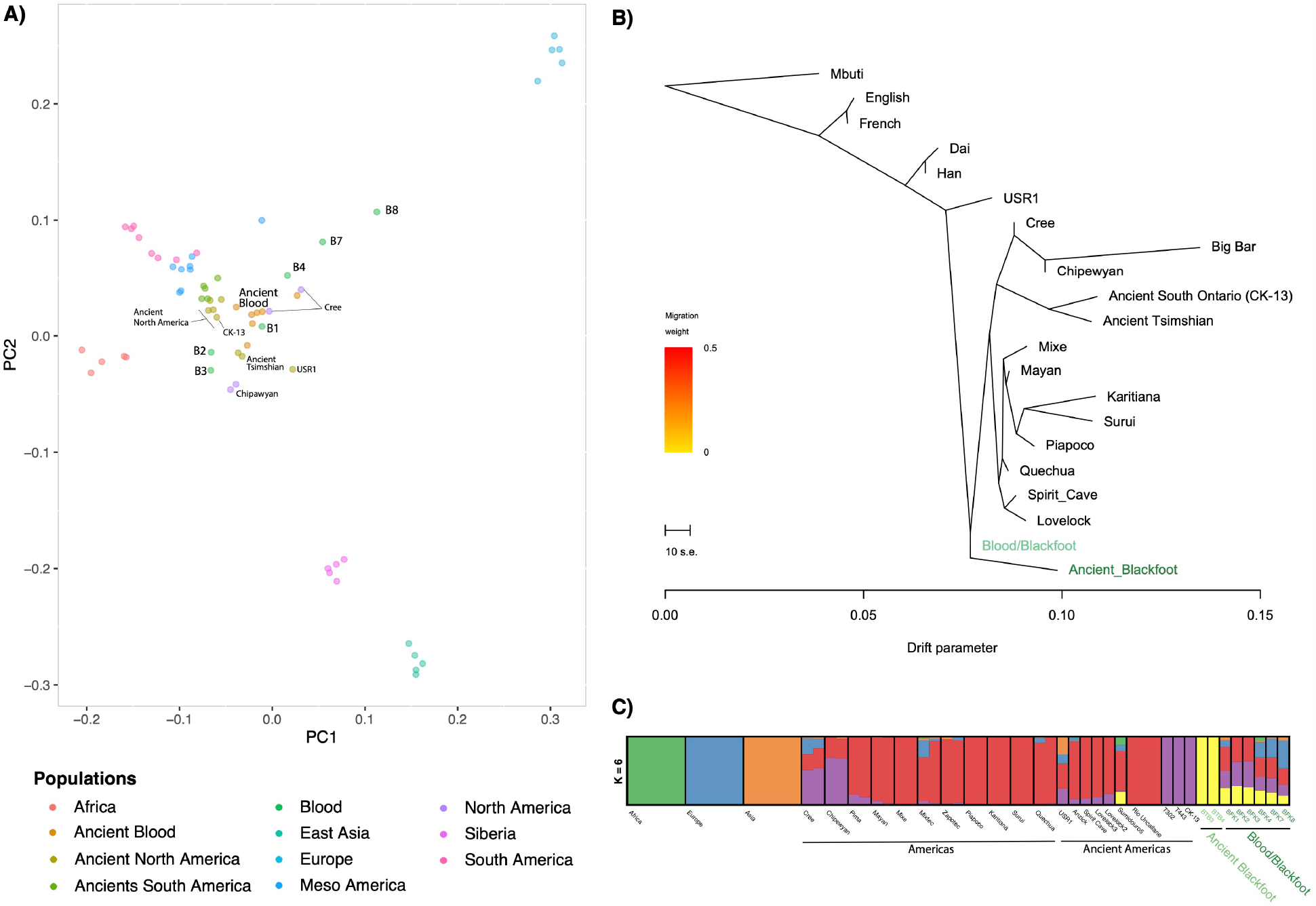
Assessing the genetic affinity of Blood/Blackfoot with the Americas. A) Principal component analysis projection of the ancient samples onto the first two principal components (PCs). B) Maximum likelihood tree generated by *TreeMix* using whole-genome sequence data from the Simons Genome Diversity Project and ancient genomes from previously published studies. C) Ancestry clusters generated with *ADMIXTURE* of modern and ancient genomes from the Americas at *K*=6 clusters, which was chosen through cross-validation.

**Figure 3.**
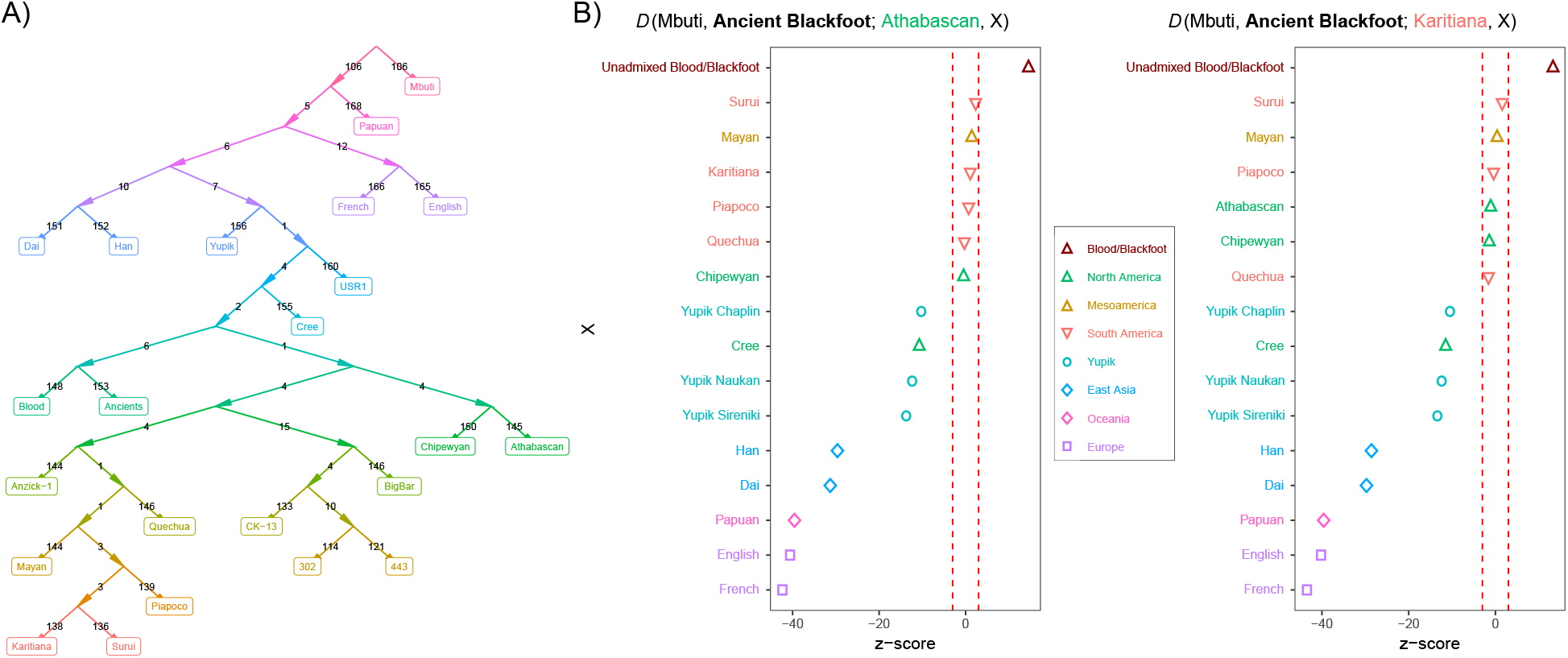
Demographic model estimated by qpGraph with no migration events and *D*-statistics. A) Graph depicts the historical Blackfoot and present-day Blood/Blackfoot forming a clade that splits with the Northern and Southern American lineages, mirroring the findings of the *TreeMix* analysis of Figure 2B. The Cree population splits more ancestral due to the individuals harboring non-negligible European ancestry components, which pushes its split deeper into the past.B) *D*-statistics showing excess allele sharing between ancient and present-day Blood/Blackfoot and ancient Blood/Blackfoot equally distant to populations in North America (ANC-B) and South America (ANC-A). Here, dashed red lines represent significance thresholds.

With multiple genomic analyses showing the ancient Blood/Blackfoot clustering together with present-day Blood/Blackfoot but on a separate lineage from other North and South American groups, we created a demographic model using momi2 (Figure 4), which utilized the site frequency spectra of present-day Blood/Blackfoot, Athabascan (as a representative of Northern Native American lineage), Karitiana (as a representative of Southern Native American lineage), and Han, English, Finnish, and French representing lineages from Eurasia. The best-fitting model shows a split time of the present-day Blood/Blackfoot at 18,104 years ago, followed by a split of Athabascan and Karitiana at 13,031 years ago (Table 2).

**Table 2.**
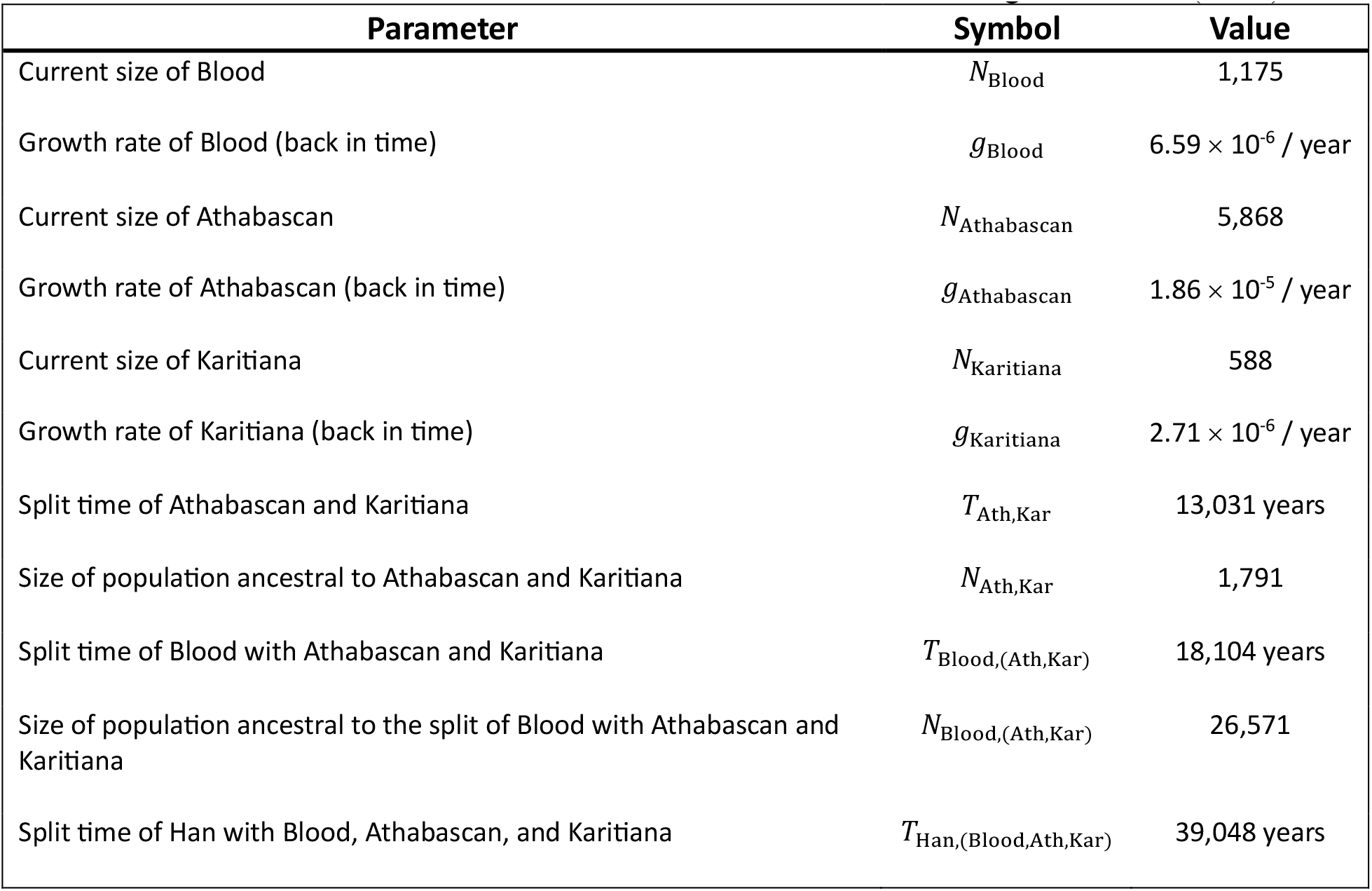
Demographic parameters inferred by momi2, relating Han, Blood, Athabascan, and Karitiana populations. All population sizes, growth rates, split times, and ancestral sizes relating the European and Han populations are fixed values from Table 2 of Jouganous et al. (2017).

| Parameter | Symbol | Value |
| --- | --- | --- |
| Current size of Blood | $N_{\text{Blood}}$ | 1,175 |
| Growth rate of Blood (back in time) | $g_{\text{Blood}}$ | $6.59 \times 10^{-6} / \text{year}$ |
| Current size of Athabascan | $N_{\text{Athabascan}}$ | 5,868 |
| Growth rate of Athabascan (back in time) | $g_{\text{Athabascan}}$ | $1.86 \times 10^{-5} / \text{year}$ |
| Current size of Karitiana | $N_{\text{Karitiana}}$ | 588 |
| Growth rate of Karitiana (back in time) | $g_{\text{Karitiana}}$ | $2.71 \times 10^{-6} / \text{year}$ |
| Split time of Athabascan and Karitiana | $T_{\text{Ath,Kar}}$ | 13,031 years |
| Size of population ancestral to Athabascan and Karitiana | $N_{\text{Ath,Kar}}$ | 1,791 |
| Split time of Blood with Athabascan and Karitiana | $T_{\text{Blood,(Ath,Kar)}}$ | 18,104 years |
| Size of population ancestral to the split of Blood with Athabascan and Karitiana | $N_{\text{Blood,(Ath,Kar)}}$ | 26,571 |
| Split time of Han with Blood, Athabascan, and Karitiana | $T_{\text{Han,(Blood,Ath,Kar)}}$ | 39,048 years |

**Figure 4.**
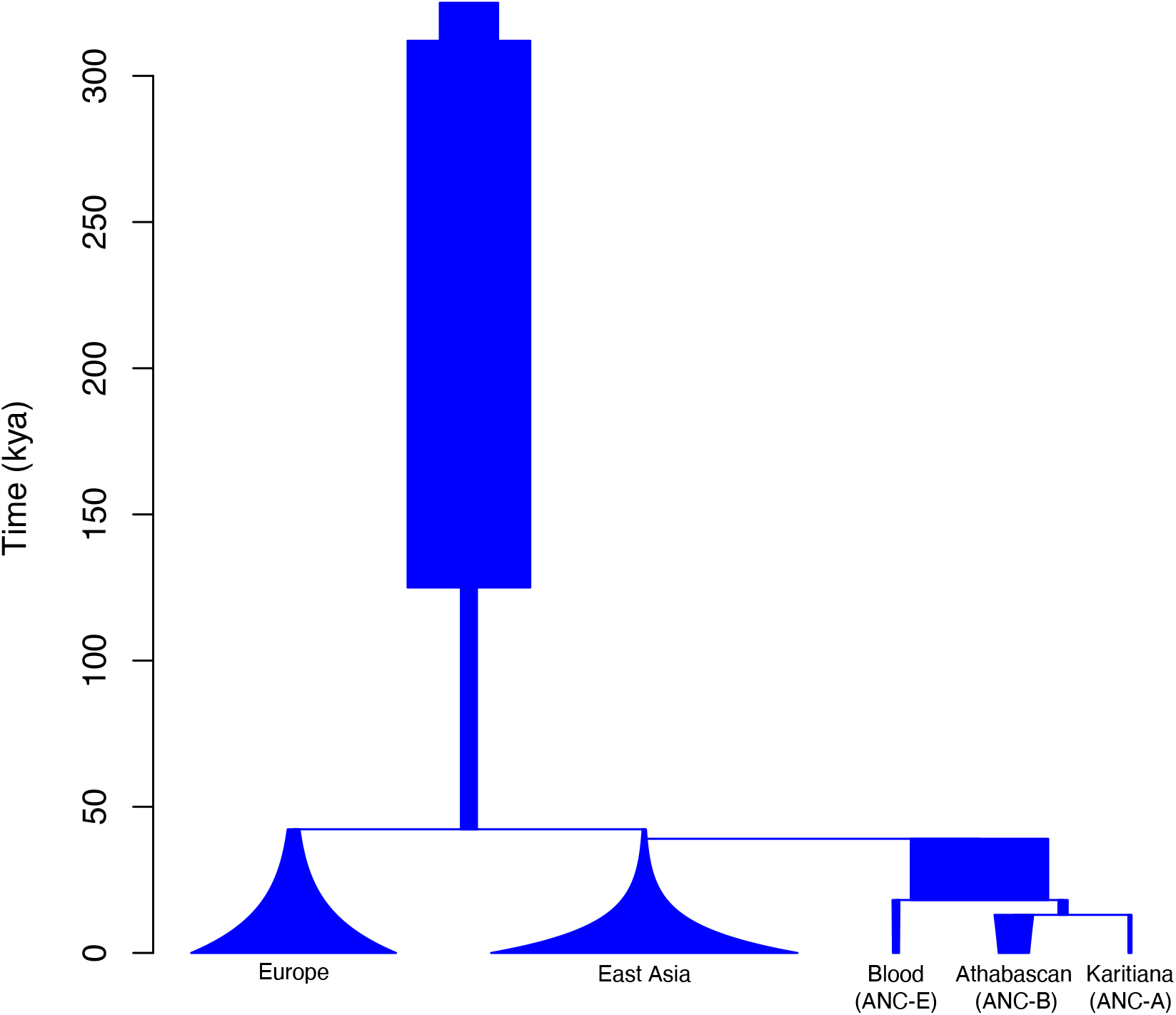
Demographic model estimated by momi2. Population sizes, growth rates, split times, and ancestral sizes relating the European and Han populations are fixed values (from Table 2 of Jouagnous et al. 2017), whereas the remaining parameter estimates can be found in Table 2 and were inferred by momi2. All times and population sizes are depicted proportional to each other.

## Discussion

In this study, we sequenced genomes of seven historic Ancestors associated with the Blackfoot Confederacy and six present-day members of the Blood (Kainai) First Nation/Blackfoot Confederacy to infer population history in combination with archaeological, oral history, and historical linguistic information. The genomic data generated was compared to populations in the Americas to infer population structure. The historic Ancestors analyzed in this study show genetic continuity with present-day members of the Blood/Blackfoot demonstrating that these genomic patterns are grounded in the traditional Blackfoot Confederacy homelands at the time of European contact.

The *TreeMix*, qpGraph, *D*-statistics and demographic model results show that the ancient Blood/Blackfoot and present-day Blood/Blackfoot individuals belong to the same lineage that split near the terminal end of the Late Pleistocene after the Ancient Beringian lineage split, but before the separation of the Northern and Southern Native American lineages. Interestingly, the Blood/Blackfoot do not group with other speakers of Algic languages, including central Algonquians. This ancient split can be compared with recent historical linguistic analyses of the Blackfoot (*16, 17*) that show the deep antiquity of the Blackfoot language within the Algic family, specifically that certain elements of Blackfoot are older than proto-Algonquian language and likely were spoken by Indigenous peoples in the aboriginal homelands of the Blackfoot Confederacy as supported by placenames and oral histories (*13, 18*). This finding changes the traditional anthropological assumption that the Blackfoot language (and, by extension, its speakers) originated in the North American Great Lakes, where Algonquian purportedly evolved (*19*). Rather, our study aligns with the Goddard (*17*) reconstruction of a west-to-east cline in the Blackfoot language and population.

A plausible scenario consistent with our multiple sources of knowledge is the presence of an ancient population in the Blackfoot homeland that emerged in the Late Pleistocene and that was severely impacted by the ashfall from Mount Mazama eruption about 7,600 years ago (*20*). Many of these ancestors might have departed the homeland until the grassland recovered and the bison returned. Perhaps linguistic elements were shared during this temporary movement, along with other cultural practices such as secondary, red ocher burial bundles and copper artifacts that were pervasive in the Great Lakes (*21*).

The finding of a previously unidentified lineage in the Americas that emerged in the Late Pleistocene builds on a pattern of multiple lineages being identified prior to the split of the Northern and Southern Native American lineages including the Ancient Beringian lineage (*2*), the population represented by the lineage associated with the Big Bar ancestor (*3*), and “ghost” populations contributing to the genomic diversity in the Americas most recently identified through statistical analyses (*5, 6*). These findings are potentially consistent with models of multiple populations radiating from a single source into vast geographic lands, evolving in relative isolation into different lineages early on, before interacting with nearby communities later in time as we infer here with the Blood/Blackfoot.

We take this opportunity of identifying a previously unknown genomic lineage to adjust practice in the field. The genomics literature currently uses names of lineages that can be confusing, especially out of the context of genomic literature. For example, “Native American” usually refers to Indigenous peoples in the United States but “Northern Native American” and “Southern Native American” are names given to genomic lineages of Indigenous peoples that spans the Americas, both within and out of the United States. To minimize perplexing terminology, we suggest following the lead of Scheib and colleagues (*22*) and expanding their terminology for lineages as in Figure 5.

**Figure 5.**
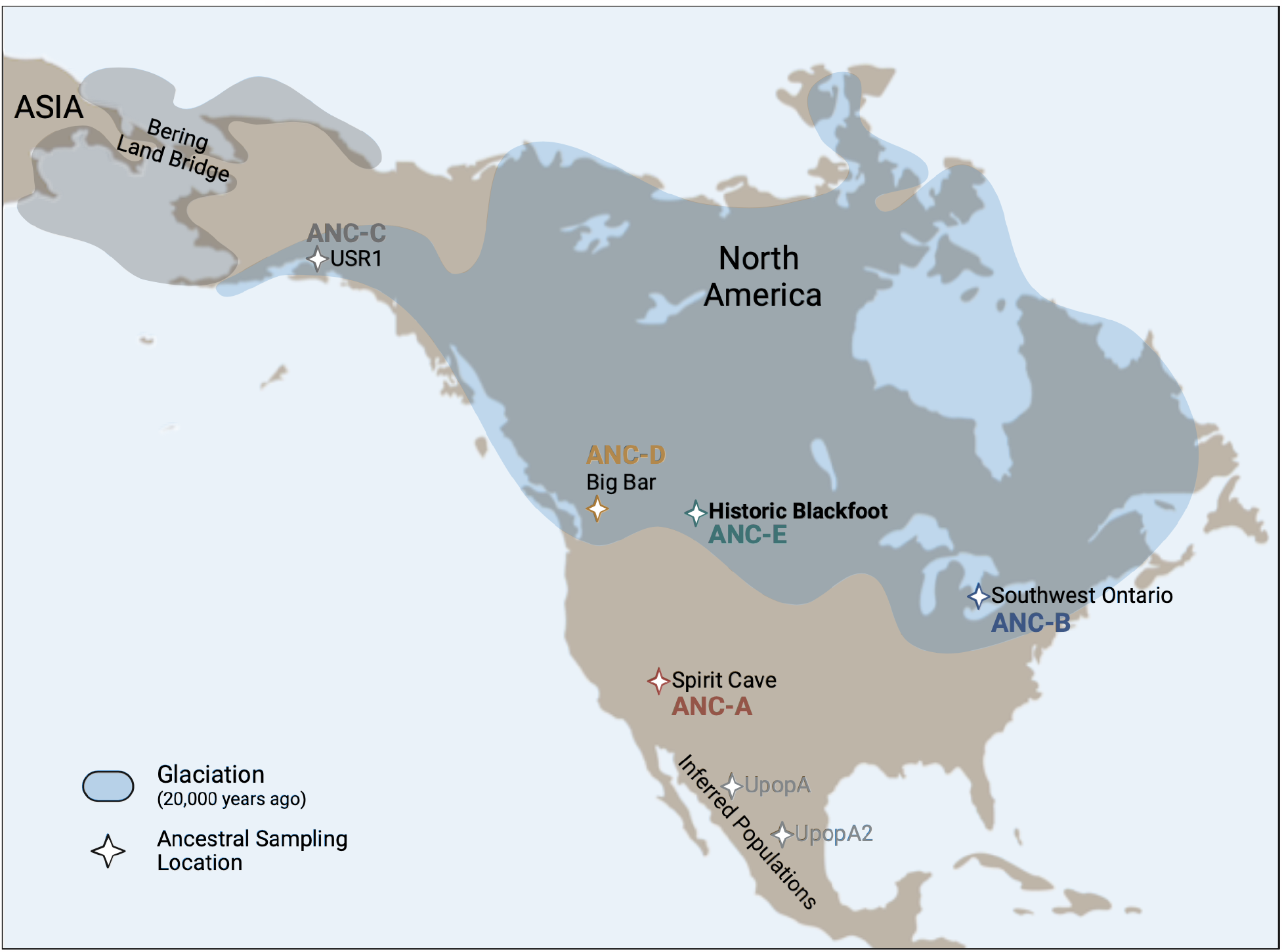
Genomic lineages of the Americas that evolved in the Late Pleistocene. Nomenclature follows from (22) with USR1 (2) representing ANC-C, Big Bar (3) representing ANC-D and Historic Blackfoot representing ANC-E.

## Materials and Methods

### Sampling and radiocarbon dates

Samples were obtained from three historical ancestors identified as Blood/Blackfoot at the Smithsonian Institution and four historical ancestors curated at the Blackfeet Tribal Historic Preservation Office (*n*=7). The Blackfoot practiced tree or scaffold burial followed by secondary burial (*10*). However, in the historic period secondary burial fell into disuse possibly due to epidemics and famine. Thus, samples from historical ancestors were not excavated but collected from the surface of old burials by various individuals (including the US Army Surgeon) in the 19^th^ and 20^th^ centuries. A few of these remains were then placed in repositories. While the exact location of these findings is uncertain, in all cases they were identified as Blood/Blackfoot based geographic location within Blackfoot lands.

Radiocarbon dating was conducted on tooth collagen at the University of Georgia Center for Applied Isotopic Studies (UGAMS). Pretreatment, collagen extraction, and isotopicmeasure-ments followed the standard protocol for this laboratory. We calibrated the dates with OxCal (*23*) and IntCal20 (*24*).

### DNA extraction, library builds, and sequencing

#### DNA extraction

DNA was extracted from the seven historical ancestors using an ancient DNA extraction protocol previously developed and optimized for ancient DNA analysis (*25*) at the Carl R. Woese Institute for Genomic Biology, University of Illinois at Urbana-Champaign. All DNA extractions were performed in a low-template DNA laboratory dedicated to the analysis of ancient DNA. To avoid external or cross-sample contamination, extraction and PCR-negative controls were included with each round of sample processing, and no more than eight samples were processed at a time.

Following the DNA extraction of samples from historical ancestors, DNA extractions from present-day community members were taken from saliva samples collected using Oragene DNA Saliva Kits and DNA extraction kits from DNA Genotek under University of Illinois IRB protocol #10538 and Blood (Kainai) First Nation Tribal Council approval. Following DNA extraction and quantification, DNA extracts were sequenced by the University of Illinois Roy J. Carver Biotechnology Center.

#### Genomic library preparation and sequencing

Genomic libraries were constructed using the NEBNext® Ultra II™ DNA Library Prep kit and NEBNext® Multiplex Oligos (Unique Dual Indexes) for Illumina® following an optimized protocol for ancient DNA analysis (*25*). Libraries were pooled and shotgun sequenced using a 100 base single-end sequencing approach on an Illumina HiSeq 6000 platform at the Roy J. Carver Sequencing Facility. Based on the sequencing of this pool of libraries, we identified two samples (BTB3 and BTB4) that had relatively higher endogenous DNA content, which were re-sequenced to higher depth of coverage using a 150-base paired-end approach on the same platform.

### Alignments, haplogroup calls, and DNA damage

#### Nuclear genome alignment, variant calls, and genomic sex estimation

The raw sequences were trimmed for Illumina adapters using AdapterRemoval2 (*26*) and aligned to the hg19 human reference sequence using the BWA mem algorithm (*27*). Genotypes for all ancient individuals were called with the ancient DNA caller ARIADNA, which employs a machine learning method to overcome issues with DNA damage and contamination (*28*). The contemporary individuals were called with bcftools (*29*), using a minimum mapping quality of 30 and a minimum base quality of 20. The resulting variant call files (VCFs) were further filtered to remove genotype calls with allele counts below three. The VCFs were then merged with modern and ancient samples from the Americas with bcftools. Genomic sex was estimated for all historic individuals (*30*). We calculated the breadth of genome coverage (i.e., the percentage of the genome that has ≥1X read coverage) and the mean depth of coverage (the mean number of reads that mapped at any location across the genome, Table 1).

#### Mitochondrial haplogroup calls

The raw sequences were also aligned to only the human mitochondrial DNA reference sequence (ChrM, GenBank accession NC_012920.1) using the same alignment parameters and programs. VCFs were generated using SNVer (*31*), and mitochondrial haplogroups were identified using mitotool (*32*). Depth and breadth of mitochondrial genome coverage was calculated for all seven historical ancestors (Table 1).

#### DNA damage estimation

We assessed whether DNA showed damage patterns that are characteristic of ancient DNA by quantifying damage for nuclear genome alignment files in MapDamage2 v. 2.0.5 (*33*) using a fragment size of 70 bases. Damage patterns (Supplementary Figure 1) were visualized using a custom R script.

### Principle component analysis, model-based clustering, trees, and D-statistics

#### SmartPCA

We conducted principal component analysis using the ‘smartpca’ program from the EIGENSOFT v7.2.1 package (*34*). Principal components (PCs) were estimated with the ‘poplistname’ option and using representative individuals from present-day Indigenous populations from Simons Genome Diversity Panel (*35*) The ancient individuals were then projected onto the computed PCs with the ‘lsqproject: YES’ option. No outliers were excluded, linkage disequilibrium pruning was not performed, and the analysis was based on 1,231,622 loci.

#### Assessment of population structure using ADMIXTURE

We started with the identical filtered dataset of called genotypes described above. We further pruned the dataset by removing sites in strong linkage disequilibrium (r^2^ > 0.1) using PLINK (*36*). The program *ADMIXTURE* (*37*) was used to assess global ancestry of the ancient and present-day samples from this study. We computed cluster membership for *K*=2 through *K*=8, running 10 replicates for each value of *K* while generating pseudo-random seed with the *-s* option (Supplementary Figure 2). The replicate with the highest likelihood was then chosen for each *K*. The PONG program (*38*) was used to visualize the admixture plots.

#### TreeMix analysis

We started with the filtered dataset of called genotypes. *TreeMix* (*39*) was applied to the dataset to generate maximum likelihood trees and admixture graphs from allele frequency data. The Mbuti from the Simons dataset was used to root the tree (with the *–root* option). We accounted for linkage disequilibrium by grouping *M* adjacent SNPs (with the *–k* option), and we chose *M* such that a dataset with *L* sites will have approximately *L/M* ≈ 20,000 independent sites. At the end of the analysis (i.e., number of migrations) we performed a global rearrangement (with the *–global* option). We considered admixture scenarios with *m* = 0 and *m* = 3 migration events. We performed 20 iterations for each admixture scenario, choosing the highest likelihood for each.

#### qpGraph analysis

We filtered the genotype calls of the Simons Genome Project (*35*) to include only Mbuti (*n*=4), Papuan (*n*=15), French (*n*=3), English (*n*=2), Dai (*n*=4), Han (*n*=3), Eskimo (*n*=5; Chaplin, Nakaun, and Sireniki), Cree (*n*=2), Chipewyan (*n*=2), Quechua (*n*=3), Mayan (*n*=2), Piapoco (*n*=2), Surui (*n*=2), and Karaitiana (*n*=3) individuals. In this manuscript, we replace the term Eskimo with Yupik for the Simons Genome Project sample data. Using the bcftools *--merge* option, we then intersected these genotype calls with those from Athabascan (*n*=2; (*40*)) individuals, the ancient Blood/Blackfoot and present-day Blood/Blackfoot individuals from this study, and the ancient Anzick-1, 302, 443, USR1, Big Bar, and CK-13 individuals (*2, 3, 22, 41, 42*). We then further filtered this dataset to include only biallelic SNPs with no missing data using the bcftools *-m2 -M2 -v, --min-ac 1:minor*, and *filter -I ‘F_MISSING <= 0*.*0’* options. Finally, to guard against artifacts due to DNA damage, we removed C/T and G/A SNPs from the final dataset, which resulted in 10,385 loci for analysis. To perform qpGraph analysis (*43*), we employed the R package ADMIXTOOLS2 (https://uqrmaie1.github.io/admixtools/index.html, ADMIXTOOLS2 is currently under preparation). Using a setting with no migration events, we embarked on a graph search with a random graph with Mbuti fixed as the outgroup, running the search for 1,000 iterations.

#### Demographic model inference with momi2

We extracted English (*n*=2), Finnish (*n*=1), French (*n*=2), Han (*n*=3), and Karitiana (*n*=3) individuals from genotype calls of the Simons Genome Diversity Project (*35*), and intersected these calls with genotype calls from Athabascan (*n*=2; (*40*)) and the unadmixed present-day Blood/Blackfoot individuals (*n*=2, BFK2 and BFK3) from this study using the bcftools *-merge* option. We then filtered this dataset to include only biallelic SNPs using the bcftools *-m2 -M2 -v* and *--min-ac 1:minor* options, which resulted in 2,221,154 loci for analysis. Using momi2 (*44*), we computed allele counts using momi.read_vcf with the *--no_aa* option and calculated the site frequency spectra with block jackknifing using mom.extract_sfs with 100 blocks.

Fixing the population sizes, growth rates, split times, and ancestral sizes relating the European (English, Finnish, and French) and Han populations with values from Table 2 of Jouganous et al. (*45*), we estimated population sizes, growth rates, split times, and ancestral sizes relating the Blood/Blackfoot, Athabascan, and Karitiana populations, as well as the split time of these populations with Han using momi2 assuming a generation time of 29 years. The relative order of population splits and their ages were constrained based on evidence from the qpGraph topology observed in Figure 3. Specifically, the split with Han had an upper bound of 42,300 years ago (fixed split time between Han and Europeans of Jouganous et al. (*45*)) and a lower bound of 12,600 years ago (age of the Anzick-1 sample), the split of Blood/Blackfoot with Athabascan had a lower bound of 12,600 years ago and an upper bound as the estimated split with Han, and the split of Athabascan with Karitiana had a lower bound of 12,600 years ago and an upper bound as the estimated split of Blood/Blackfoot with Athabascan. Optimal model parameters were obtained using the truncated Newton algorithm.

#### D-statistic tests

We used the POPSTATS Python program (*46*) to assess whether the Blood/Blackfoot represent a genomic lineage distinct from ANC-A and ANC-B. Specifically, we ran *D*-statistic tests in the form of *D* (Mbuti, TestPop; B, X), in which ‘X’ was all present-day, non-African populations from the Simons Genome Diversity Project (*35*) used in the qpGraph analysis, while ‘TestPop’ was each one of three subsets of Blood/Blackfoot individuals, namely, (a) ancient Blackfoot, (b) unadmixed Blood/Blackfoot, and (c) admixed Blood/Blackfoot. As baselines for these tests (denoted by ‘B’ in the *D*-statistic test form), we selected the Athabascans and Karitiana populations as they are representative of the ANC-B and ANC-A lineages, respectively. In these tests, a significant positive z-score (> 3) suggest excess allele sharing between ‘TestPop’ and ‘X’ in relation to ‘B’, whereas a significant negative z-score (< -3) denotes excess allele sharing between ‘TestPop’ and ‘B’ in relation to ‘X’. Values for z-score between -3 and 3 mean that ‘TestPop’ is equally distant to ‘B’ and ‘X’. No pruning for linkage disequilibrium was applied. The number of polymorphic sites used in this analysis depends on the coverage of the four populations that are being compared. The minimum number of sites analyzed was 218,115 sites in *D* (Mbuti, Ancient Blackfoot; Karitiana, Surui) and the maximum was 453,046 sites in *D* (Mbuti, Admixed Blood/Blackfoot; Karitiana, Surui).

## Acknowledgments

We thank the Blood (Kainai) First Nation, Blackfoot Tribal Historic Preservation Office and Blackfoot Confederacy for support of this project. Alvaro Hernandez and Chris Wright at the Carver Biotechnology Center, University of Illinois.

## Author contributions

Conceptualization: DFF, ACEW, MNZ, RSM

Community engagement: DFF, ACEW, JM, MNZ, RSM

Data generation: AdF, FL

Data analysis: AdF, FL, MD, JL, ALCdS

Funding acquisition: MNZ, RSM

Project administration: MNZ, RSM

Writing: DFF, ACEW, JM, AdF, FL, MD, JL, MNZ, RSM, ALCdS

## Data and materials availability

The Blackfoot Confederacy will review requests for genomic data before access can be granted. Please send requests to

## Supplementary Materials

Figs. S1 to S4

Tables S1 to S2

References (*2,3,35,42,47,48*)

