## Supplemental Information for "Genomic analyses correspond with deep persistence of peoples of Blackfoot Confederacy from glacial times"

**The PDF file includes:**

Figs. S1 to S4
Tables S1 to S2
References (2,3,35,42,47,48)

**Fig. S1. DNA damage patterns.** Historic DNA samples showed damage patterns characteristic of ancient DNA (e.g., deamination of cytosine to uracil; Hofreiter et al., 2001). A fragment misincorporation plot is shown for two individuals, BTB3 in the top panel, and BTB 4 in the lower panel. Graphs on the left and right indicate C to T and G to A transitions, respectively. The *y*-axis denotes frequency of nucleotide change from the reference sequence, and the *x*-axis denotes position along the DNA fragment with the left side of the *x*-axis showing nucleotides from the 5' end of the read going into the read from left to right, and the right side of the *x*-axis showing nucleotides from the 3' end of the read going into the read from right to left.

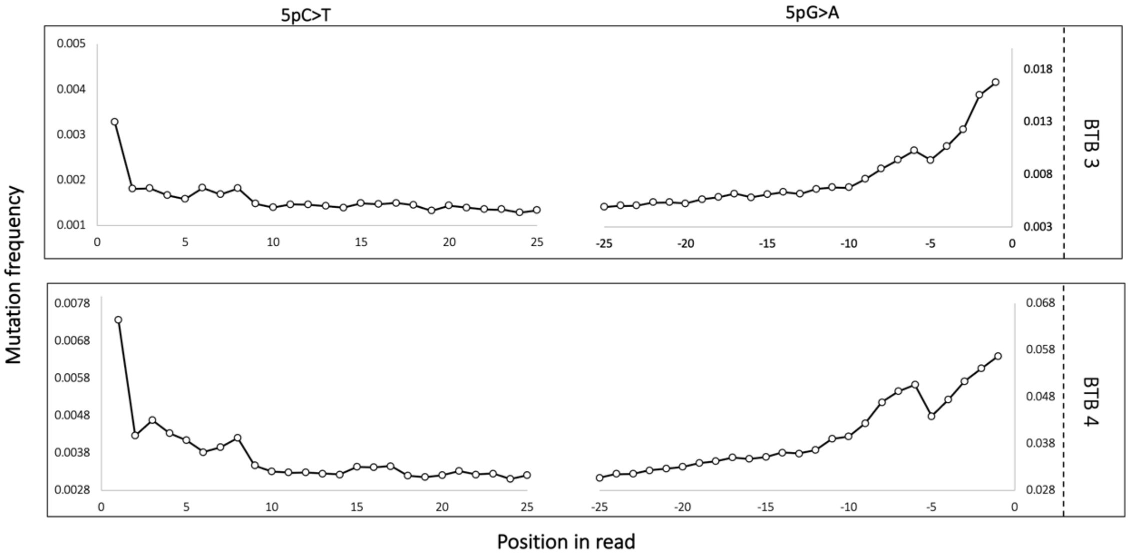

**Fig. S2. Principal Components Analysis.** Plots of PC2&3 and PC3&4.

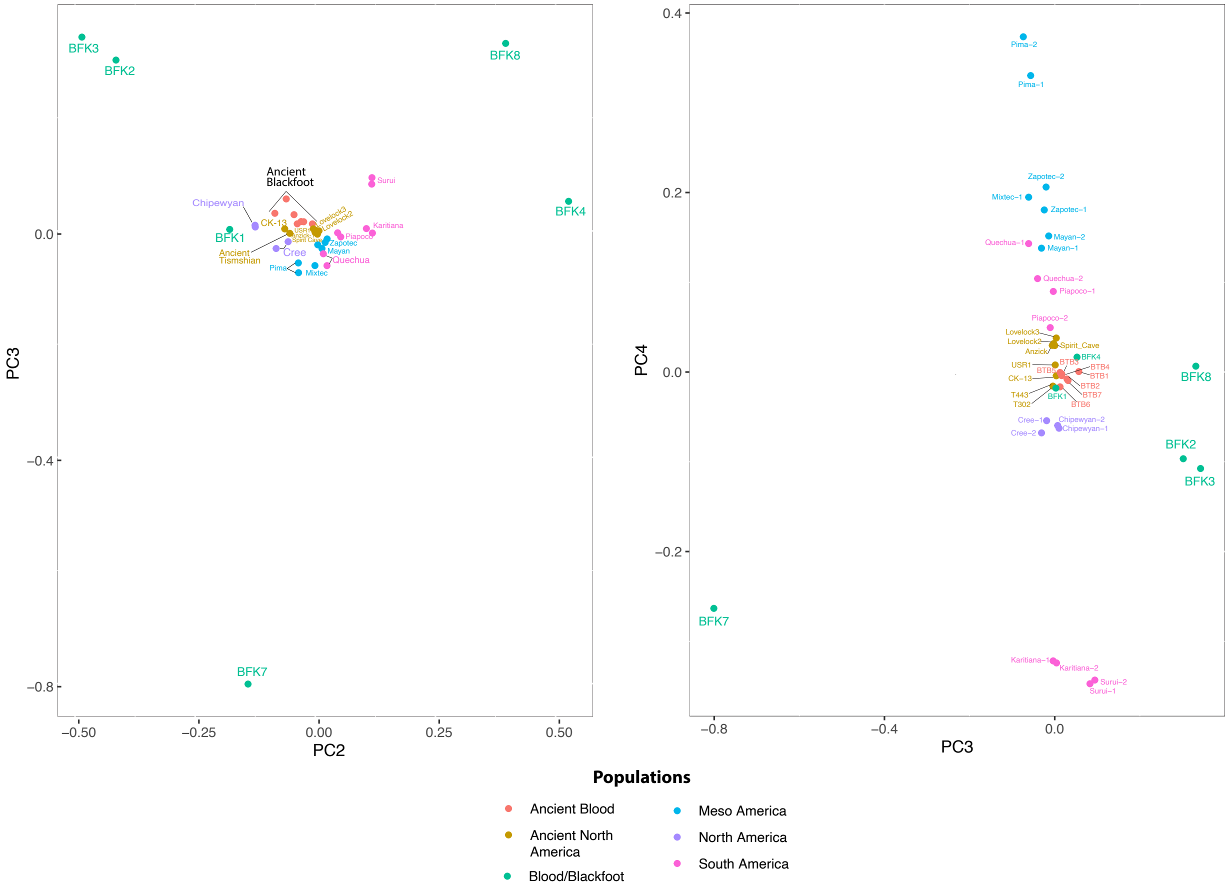


5

**Fig. S3. ADMIXTURE analysis for *K*=2 through *K*=8 clusters.** Bars represent individuals, whereas each color represents a distinct ancestral component, with bar height representing the proportion of that component comprising a given individual.

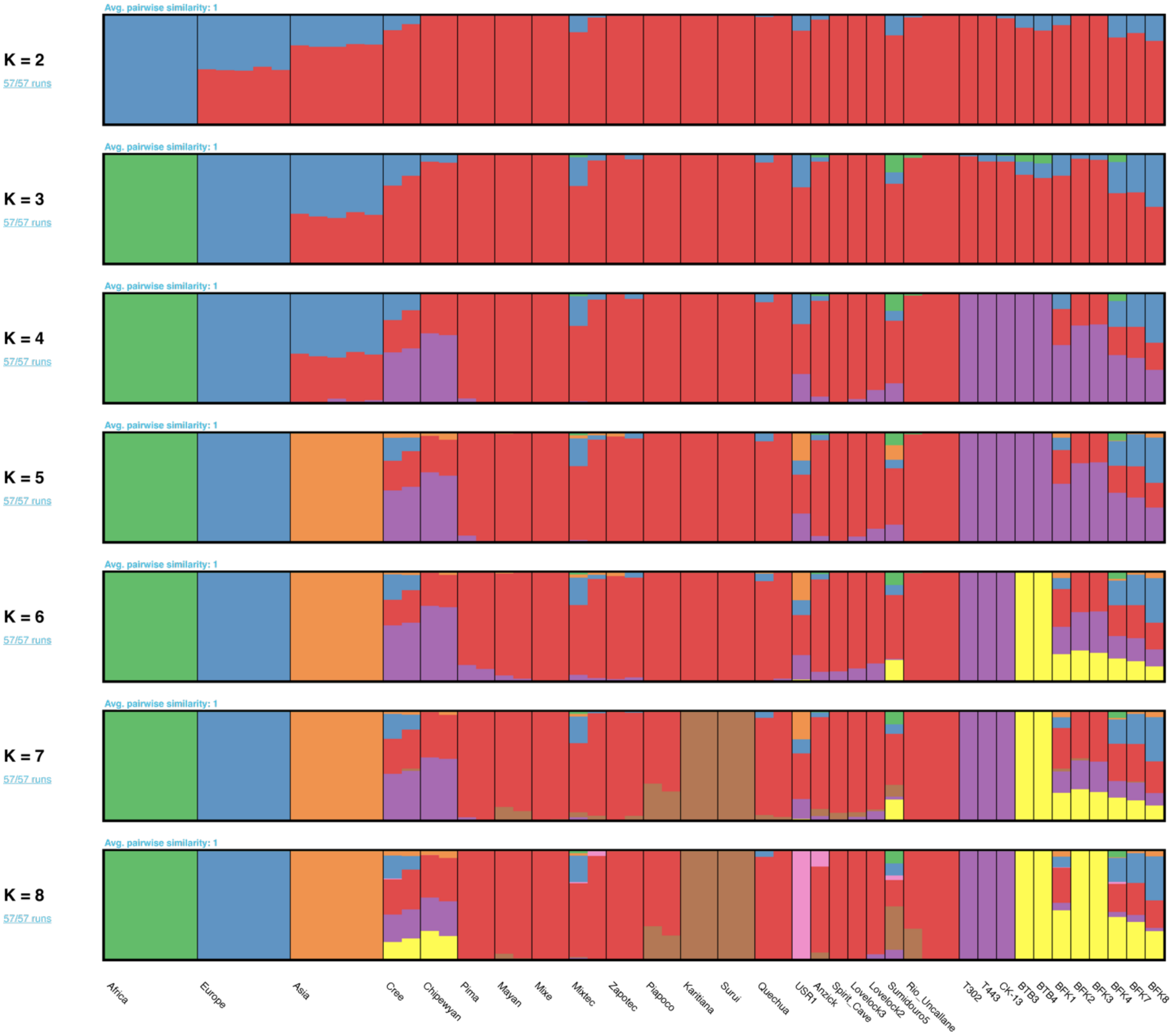


**Fig. S4. Additional *D*-statistic tests for affinity of present-day Blood/Blackfoot with other present-day populations.** A) Blood/Blackfoot genomes are equally distant to populations in

North America (ANC-B) and South America (ANC-A). B) Admixed Blood/Blackfoot showing excess allele sharing with Unadmixed Blood/Blackfoot. Dashed red lines represent significance thresholds.

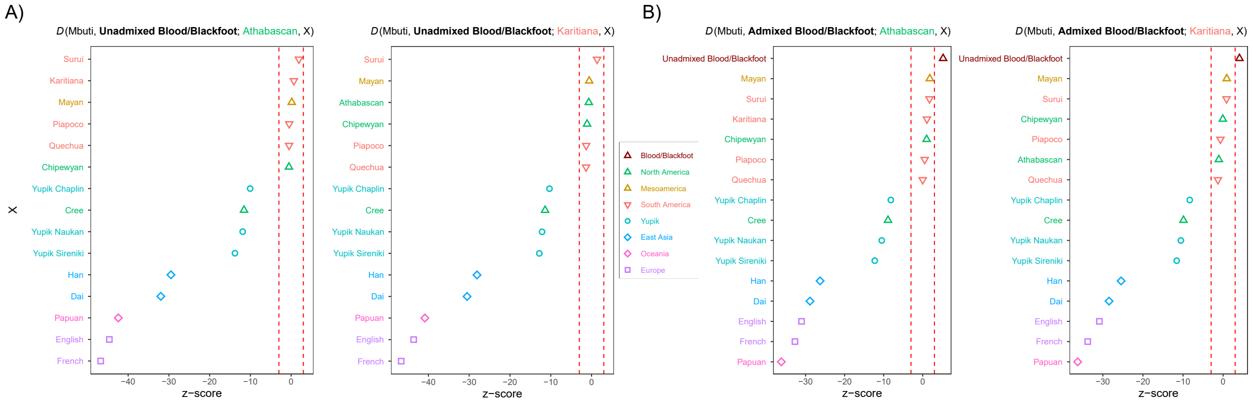

**Table S1: Ancient and contemporary samples used in this study.**

| Sample ID | Location | Average Sequence Depth | Radiocarbon Dates (BP) | Reference | ADMIXTURE | PCA |
|---|---|---|---|---|---|---|
| **Ancient Samples** | | | | | | |
| USR1 | Alaska, USA | 17 | 11.5 kyr | 2 | X | X |
| Spirit Cave | Nevada, USA | 18 | 10.7 kyr | 3 | X | X |
| Lovelock2 | Nevada, USA | 15.1 | 1.9 kyr | | X | X |
| Lovelock3 | Nevada, USA | 18.7 | 0.7 kyr | | X | X |
| Sumidouro5 | Lagoa Santa, Brazil | 15.1 | >10 kyr | | X | X |
| IL2 | Ilave, Peru | 6.5 | ~1.8 kyr | 47 | X | X |
| IL3 | Ilave, Peru | 5.9 | 1.8 kyr | | X | X |
| IL7 | Ilave, Peru | 5.6 | 1.8 kyr | | X | X |
| Anzick-1 | Montana, USA | 14.4 | 12.8 kyr | 42 | X | X |
| **Contemporary Samples (Simons Genomes Diversity Project)** | | | | | | |
| S_Pima-1 | Mexico | 43.03 | | 35 | X | X |
| S_Pima-2 | Mexico | 37.84 | | | X | X |
| S_Mixtec-1 | Mexico | 44.48 | | | X | X |
| S_Mixtec-2 | Mexico | 46.91 | | | X | X |
| B_Mixe-1 | Mexico | 40.13 | | | | X |
| S_Mixe-2 | Mexico | 46.94 | | | X | X |
| S_Zapotec-1 | Mexico | 45.75 | | | X | X |
| S_Zapotec-2 | Mexico | 39.23 | | | X | X |
| S_Mayan-1 | Mexico | 37.77 | | | X | X |
| S_Mayan-2 | Mexico | 45.13 | | | X | X |
| S_Piapoco-1 | Colombia | 40.88 | | | X | X |
| S_Piapoco-2 | Colombia | 41.01 | | | X | X |
| S_Quechua-1 | Peru | 42.04 | | | X | X |
| S_Quechua-2 | Peru | 44.82 | | | X | X |
| S_Quechua-3 | Peru | 59.78 | | | X | X |
| S_Karitiana_1 | Brazil | 44.93 | | | X | X |
| S_Karitiana_2 | Brazil | 39.49 | | | X | X |
| B_Karitiana_3 | Brazil | 37.39 | | | X | X |
| S_Surui_1 | Brazil | 41.5 | | | X | X |
| S_Surui_2 | Brazil | 35.63 | | | X | X |
| S_Eskimo_Chaplin-1 | Russia | 45.64 | | | X | X |
| S_Eskimo_Naukan-1 | Russia | 49.09 | | | X | X |
| S_Eskimo_Naukan-2 | Russia | 46.31 | | | X | X |
| S_Eskimo_Sireniki-1 | Russia | 43.61 | | | X | X |
| S_Eskimo_Sireniki-2 | Russia | 49.37 | | | X | X |
| S_Chipewyan-1 | Canada | 40.33 | | | X | X |
| S_Chipewyan-2 | Canada | 42.94 | | | X | X |
| S_Cree-1 | Canada | 36.69 | | | X | X |
| S_Cree-2 | Canada | 40.42 | | | X | X |
| S_Dai-1 | China | 49.51 | | | X | X |
| S_Dai-2 | China | 39.75 | | | X | X |
| S_Dai-3 | China | 39.05 | | | X | X |
| S_Han-1 | China | 68.83 | | | X | X |
| S_Han-2 | China | 42.36 | | | X | X |
| B_Mbuti-4 | Congo | 39.59 | | | X | X |
| S_Mbuti-1 | Congo | 41.77 | | | X | X |
| S_Mbuti-2 | Congo | 38.3 | | | X | X |
| S_Mbuti-3 | Congo | 38.78 | | | X | X |
| S_English-1 | England | 41.37 | | | X | X |
| S_English-2 | England | 45.54 | | | X | X |
| S_Finnish-1 | Finland | 38.85 | | | X | X |
| S_French-1 | France | 44.26 | | | X | X |
| S_French-2 | France | 40.2 | | | X | X |
| **Additional Samples** | | | | | | |
| Athabascan_1 | Canada | 23.2 | | 48 | X | X |

| | | | | | | |
|---|---|---|---|---|---|---|
| Athabascan_2 | Canada | 22 | | 48 | X | X |

**BP** = Before Present

**Table S2:** Radiocarbon results.

| Date | Ancestor | Material | Age ($^{14}C$ BP) | Age (cal yr AD) | $\delta^{13}C$ (‰) | $\delta^{15}N$ (‰) | C/N |
|---|---|---|---|---|---|---|---|
| UGAMS-36253 | BTB3 | molar (collagen) | 97 ± 19 | 1694-1726, 1811-1918 | -17.93 | 13.11 | 3.20 |
| UGAMS-36254 | BTB4 | incisor (collagen) | 96 ± 19 | 1694-1726, 1811-1918 | -18.15 | 13.60 | 3.21 |
| UGAMS-36255 | BTB5 | incisor (collagen) | 110 ± 19 | 1690-1729, 1808-1922 | -17.61 | 13.44 | 3.22 |
| UGAMS-36256 | BTB6 | incisor (collagen) | 95 ± 19 | 1694-1726, 1811-1918 | -17.85 | 13.46 | 3.19 |
